# CannSelect: A High-Quality Genotyping Platform for *Cannabis sativa*

**DOI:** 10.64898/2026.08.18.745408

**Authors:** Dustin G. Wilkerson, George M. Stack, Craig H. Carlson, Michael A. Quade, Caroline A. Dowling, Jacob A. Toth, Maylin J. Murdock, Jean Jasinski, Zachary J. Stansell, John K. McKay, Lawrence B. Smart

## Abstract

The field of genomics has enabled extraordinary progress in horticultural crop research. However, there is still a need for cost-effective, high-resolution technologies flexible to the diversity found in emerging crops. To this end, we introduce CannSelect, a high-quality genotyping platform for *Cannabis sativa.* Designed for use in diversity analyses and trait mapping, probe targets were selected from four genotyped diversity panels and a curated gene list. This platform has been used to effectively map day-neutrality in a segregating population to the *Autoflower1* locus with average capture efficiencies of 88.5%. With broad genome coverage, demonstrated target specificity, and reproducibility, CannSelect is expected to perform well across the diversity of *C. sativa.* We describe the methodology used to design CannSelect v1.0 and performance metrics for testing capture efficiency and target alignment in diverse genome assemblies. The CannSelect platform represents a robust and scalable, genome-wide genotyping tool for *C. sativa* researchers and breeders.

## Introduction

Genome sequencing studies have played a pivotal role in improving our knowledge of horticultural crops, especially in non-model species like *Cannabis sativa* L. These studies generally fit into three categories, whole-genome, whole-exome, and target-enrichment. Each with their benefits, target-enrichment offers the greatest genotyping accuracy while being the most cost-effective and adaptable to multiple versions or iterations [1,2]. Target enrichment probe sets are available for many plant species, such as peanut (*Arachis hypogaea)* [3], barley (*Hordeum vulgare*) [4], apple (*Malus domestica*) [5], and even angiosperms broadly [6] often for diversity analyses and trait mapping. In *C. sativa*, a target enrichment probe set flexible enough to provide high quality genotyping regardless of genetic background has yet to be developed.

Many recent advances have been made in *C. sativa* genomics, including diversity analyses using whole genome sequencing [7–9] and the publication of high-quality genome assemblies including the *Cannabis* pangenome [10–12]. Here, we introduce CannSelect, an Agilent SureSelect custom target enrichment probe set for *C. sativa*. We describe both the design process and a proof-of-concept analysis to map day-neutral flowering in a segregating population. Day-neutral flowering was chosen for probe set validation as it is a well understood phenotype with at least two causal loci, *Autoflower1,* identified on chromosome 1 between 17.7 and 22.9 Mb [13], and *Autoflower2,* on chromosome 8 between 64.1 and 64.6 Mb [14]. Following the initial test, changes were made to the design to optimize its capture efficiency in subsequent testing. Lastly, we demonstrate the applicability of the design in diverse backgrounds through *in silico* analysis of probe specificity across diverse genome assemblies and diversity groups.

## Results

### Probe Design

The initial intention of CannSelect was to establish a platform that would be effective across diverse genetic backgrounds and provide sufficient coverage of the *C. sativa* genome for trait mapping and diversity analyses. To accomplish this, four independent datasets from multiple genetic backgrounds were used to source polymorphic markers. These included low-pass resequencing and genotyping-by-sequencing data. Collectively, these datasets provided a total of 23.1 million genetic markers representative of the diversity within the species.

The markers in each dataset were filtered independently for call rate, percent heterozygosity, and thinned to prevent probe overlap. The datasets were then merged based on site position and proximity in the CBDRx (cs10) genome assembly [15], the reference genome at the time the probe set was designed. After another round of thinning based on marker density, a total of 579,728 polymorphic sites remained. In addition to designing probes from genome-wide, marker-based targets, a list of 148 potential gene targets was also included that contained Y-chromosome specific sequences, cannabidiolic acid synthase (CBDAS), additional cannabinoid synthases, and other agronomic and biologically-relevant genes. Agilent SureSelect probes are 120 bp in length and, following Agilent protocols, were 2X tiled over both gene and marker-based targets. Tiled probe sequences for each target were aligned back to the CBDRx (cs10) genome assembly and checked for predicted specificity, how well the probe aligns to its intended sequence.

The two versions of the CannSelect probe set described here, CannSelect v0 and CannSelect v1.0, can be differentiated by how their probe specificity was verified. The reason there are two versions discussed here is that CannSelect v0 was tested on over 1000 samples and suffered from low design specificity resulting in a large number of off-target alignments (i.e. low capture efficiency). This motivated us to modify our filtering approach in redesigning the probe set to the current CannSelect v1.0. For CannSelect v0, checking for probe specificity was based on percent identity and alignment length of a given probe’s blast hits. This left a potential 156,734 marker-based targets and 136 gene-based targets. More stringently, CannSelect v1.0 was designed using a formula for specificity that combined an alignment’s length, mismatches, and gap size which left a possible 104,314 potential marker-based targets and the final list of 130 gene targets (Supplemental Table 1). The remaining marker-based probes underwent a final round of selection focused on maximizing genome coverage while reducing the overall design size. This left CannSelect v0 with a total of 19,452 unique probes with an average probe count per megabase of 22.9 and a maximum gap size of 1.18 Mb (Supplemental Table 2). CannSelect v1.0 is a slightly smaller design with 18,040 unique probes with a mean probe count per megabase of 21.3 and a maximum gap size of 1.31 Mb (Table 1).

**Table 1:**
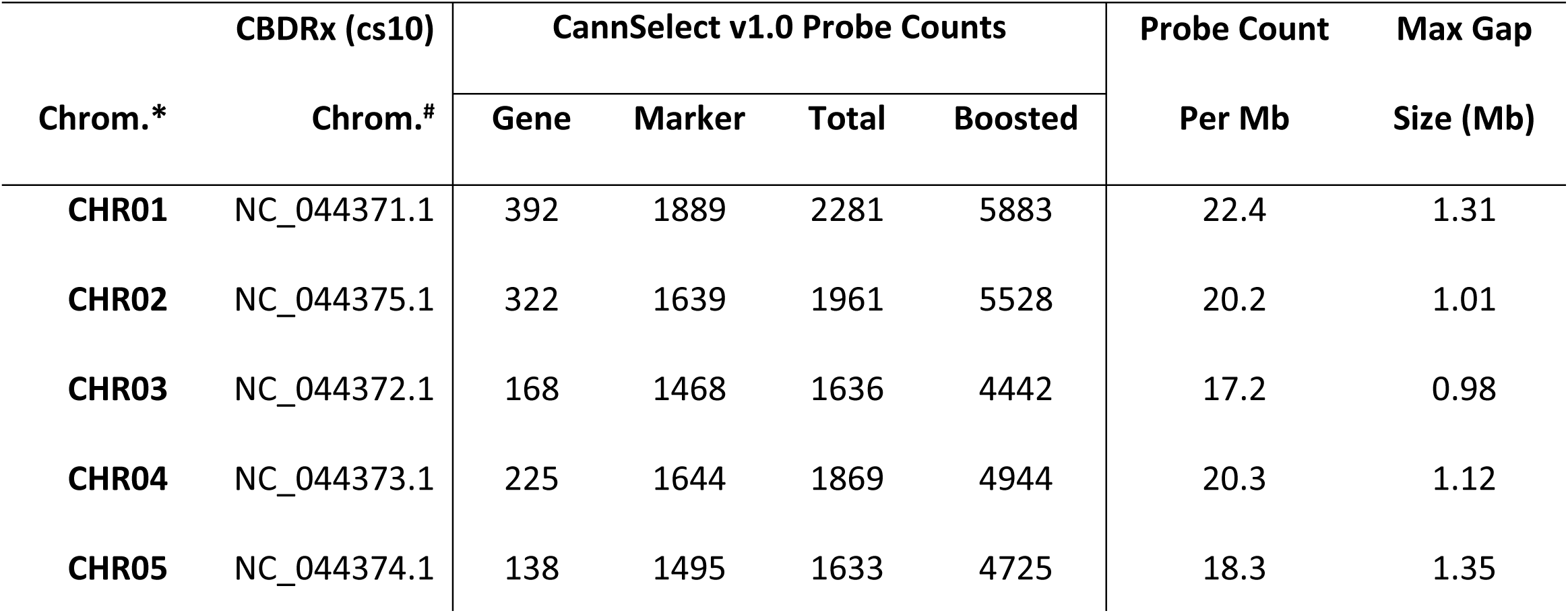
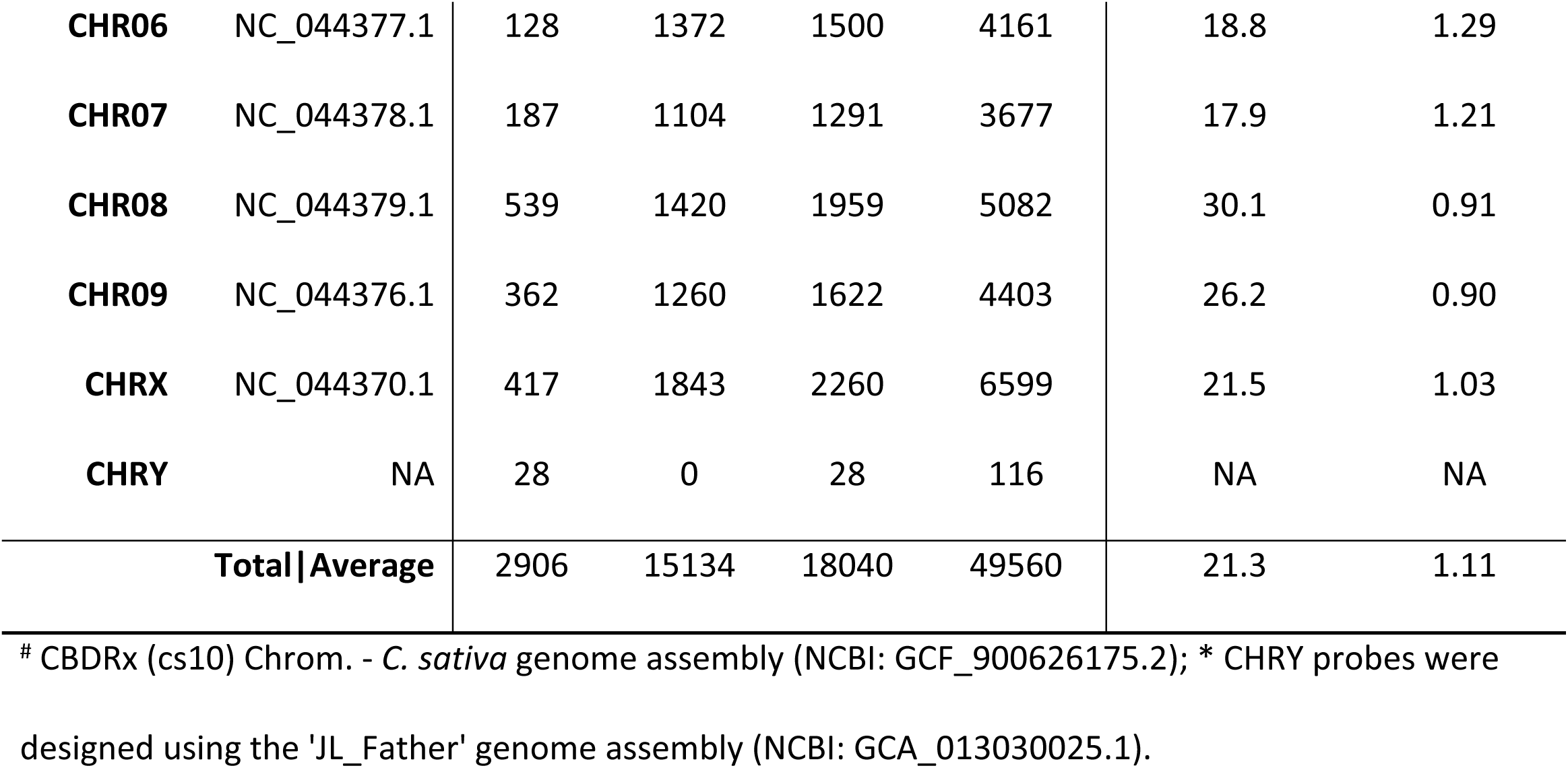
CannSelect v1.0 Probe Set Design Statistics by Chromosome.

The representation of each probe within the final design was “boosted” based on its GC content by uploading and processing it using Agilent’s SureDesign software [16]. Boosting ensures equal hybridization across all probes. This resulted in 49,560 probes for CannSelect v1.0. The final design for CannSelect v1.0 achieved a fairly uniform probe distribution across the CBDRx (cs10) assembly (Fig. 1). While only 6.3% of the 859 1 Mb intervals in the reference genome had five or fewer probes, 72.8% were covered by 10 to 50.

**Figure 1.**
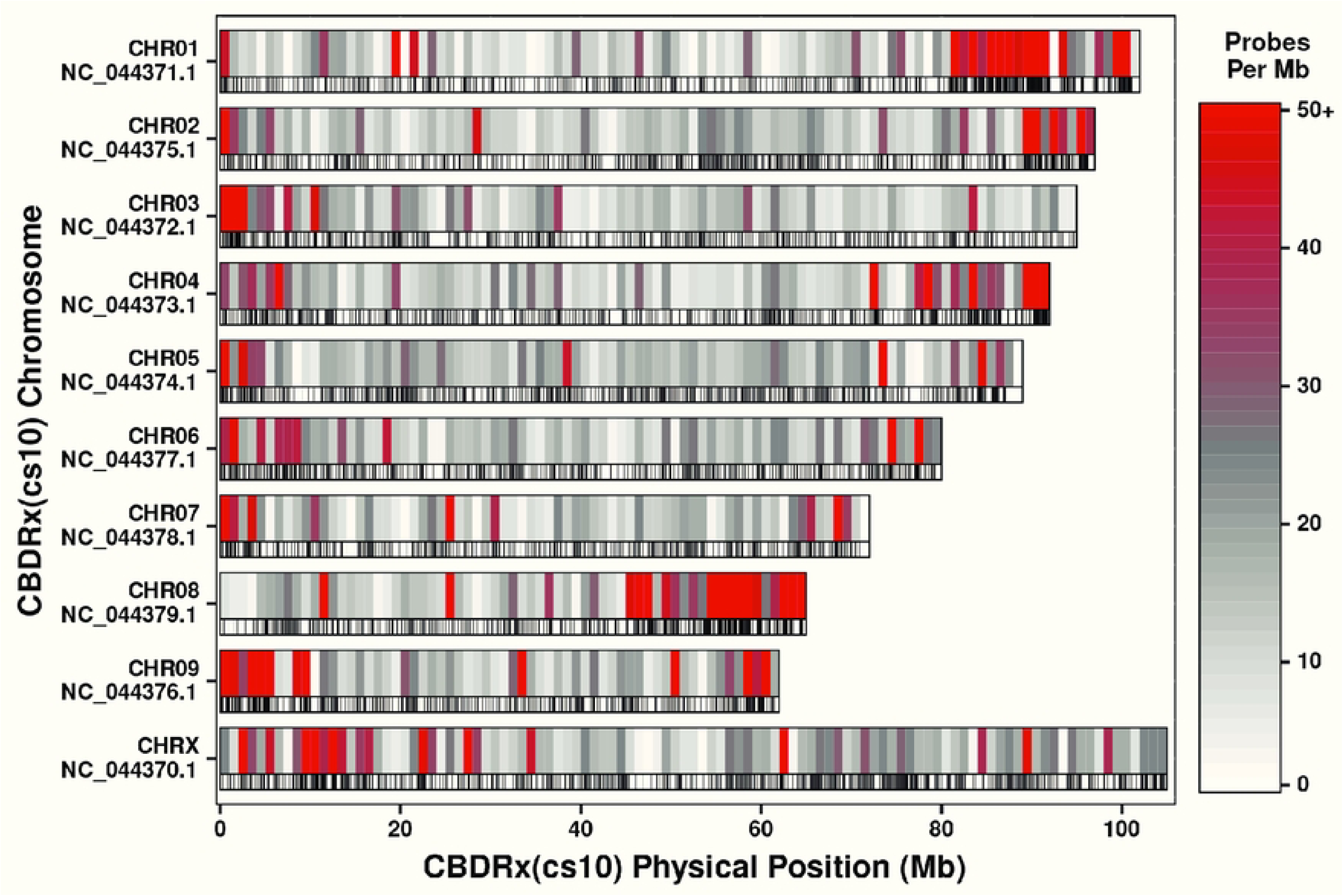
CannSelect v1.0 probe distribution and density per Mb by chromosome when aligned to CBDRx (cs10). For each chromosome, probe density is reflected in two tracks. The top track shows binned probe counts per Mb ranging from 0 (white), 25 (gray), and 50+ (red). The bottom track shows the location of each probe as a tick mark.

### Testing the CannSelect Probe Set

The CannSelect v0 probe set was used to genotype a segregating breeding population consisting of individuals of ‘Carolina Dream’ (N = 274) and ‘BaOx’ (N = 47). The population was phenotyped for day-neutral flowering, among other traits. Per sample statistics for read depth showed sufficient coverage of design targets at the chromosome level, averaging between 29.7 and 38.5X coverage (Figure 2A). At the sample level, there was a mean of 2.44 million total reads, an average of 2.27 million (93%) aligned reads after removing duplicates (Figure 2B). Across all samples, the mean percent selected was 35.8% with a standard error of 0.24% while the percentage of targets with zero coverage was just 1.75%. Regardless, the design was able to differentiate ‘Carolina Dream’ and ‘BaOx’ through multi-dimensional scaling using only on-target SNPs, with dimension 1 explaining 40.5% of the total variation and dimensions 2 and 3 explaining 15.5% and 11.4%, respectively (Fig. 2C).

**Figure 2.**
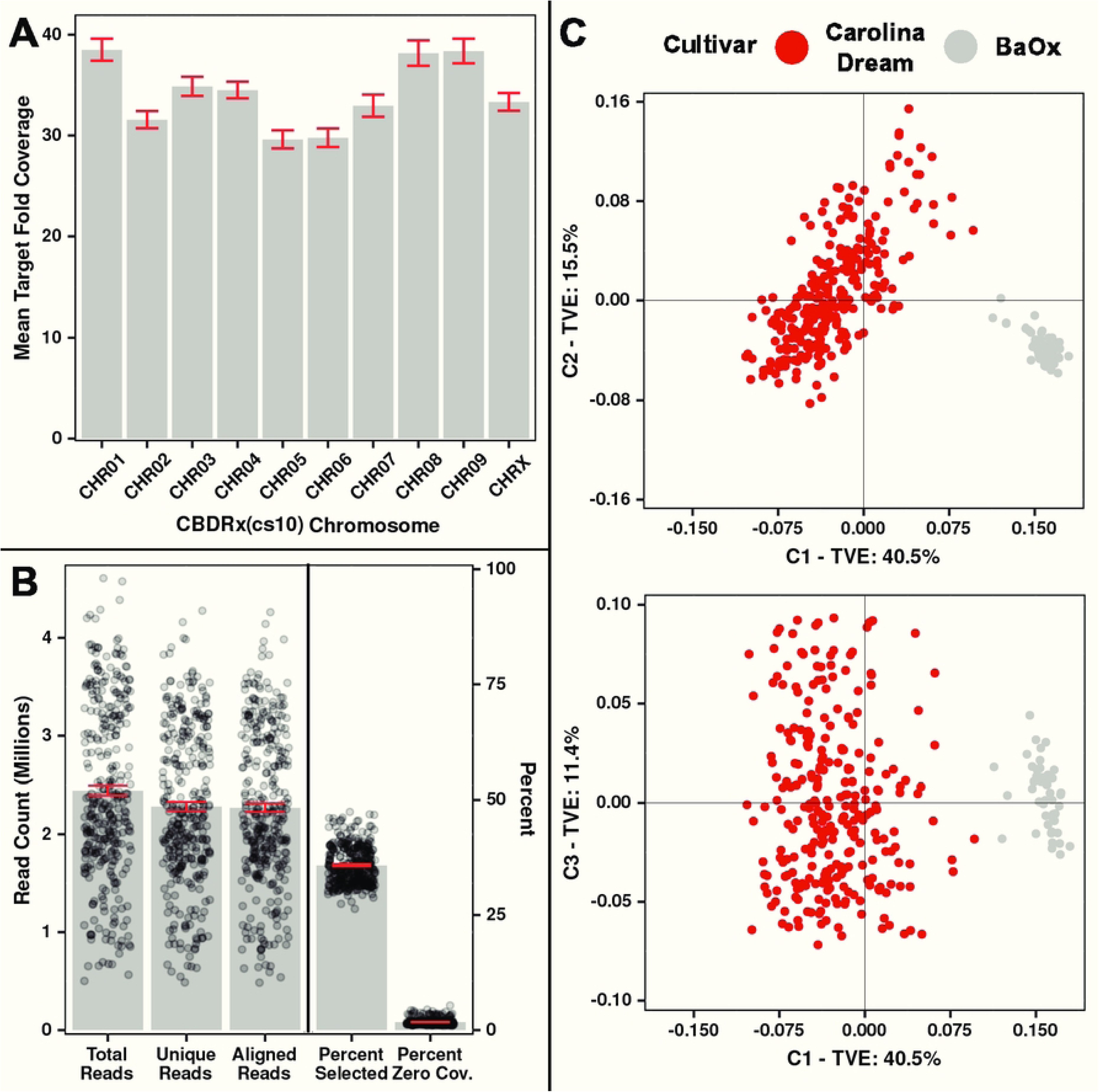
Read and probe capture statistics and MDS using CannSelect v0. A: Mean target fold coverage per chromosome as aligned to the CBDRx (cs10) genome assembly. B: Total, unique, and aligned reads (left axis) side-by-side with percent selected and percent zero coverage (right axis) for each sample. For panels A and B, the grey bars indicate the mean, while the red error bars are the standard error of the mean. C: Multi-dimensional scaling plots for 321 ‘Carolina Dream’ and ‘BaOx’ individuals. ‘Carolina Dream’ is shown in red while ‘BaOx’ is in grey. TVE: Total Variation Explained.

The 194,146 on-target SNPs were filtered for call rate, heterozygosity, and minor allele frequency prior to a GWAS of the day-neutral flowering phenotype. After filtering, missing data among the remaining 27,824 SNPs was imputed with an estimated 95.6% accuracy using LinkImputeR [17]. Repeating thresholds for call rate, heterozygosity, and minor allele frequency after imputation left 24,918 high quality SNP markers for the GWAS. Marker-trait associations for day-neutral flowering were identified using a mixed linear model with the first five principal components from a PCA and a kinship matrix to capture population structure (Fig. 3). This resulted in a significant peak on chromosome 1 at 22.47 Mb and a 1.5 LOD support interval ranging from 19.2 - 23.56 Mb as expected for the *Autoflower1* locus [13].

**Figure 3.**
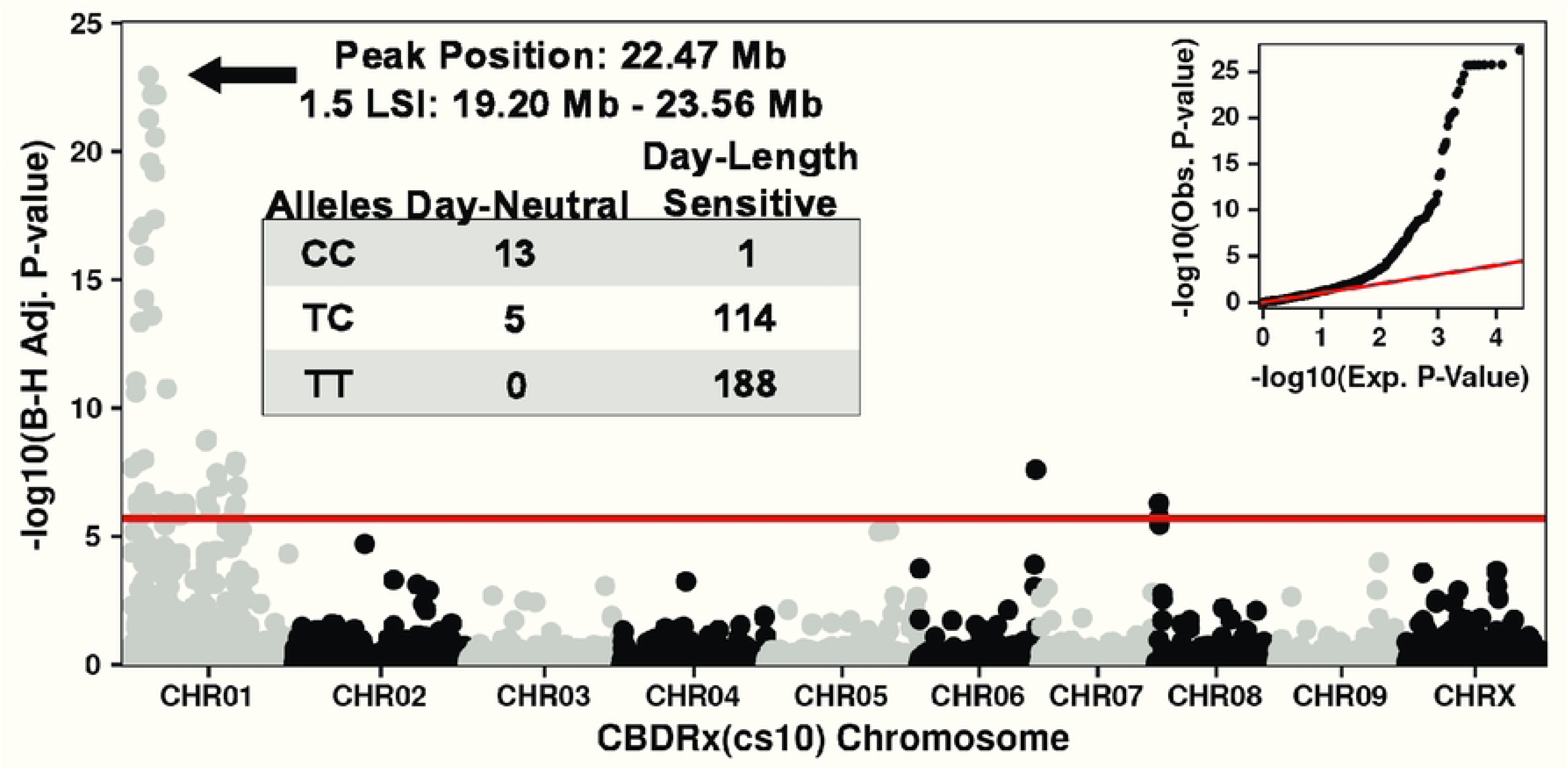
GWAS Manhattan and QQ plots mapping day-neutral flowering in the ‘Carolina Dream’ and ‘BaOx’ breeding population. The red line shows the Bonferroni threshold of - log10(0.05/24918) = 5.70. LSI: LOD Support Interval.

The design improvements made for CannSelect v1.0 resulted in substantially improved capture efficiency. Tested on 96 replicated samples of 15 diverse hemp individuals from the USDA-ARS germplasm repository and commercial checks, the mean percent selected across all samples was 88.5% with a standard error of 0.004% (Fig. 4). As each individual was represented by five plants treated as replicates, we also quantified within-individual consistency. Of the 15 individuals, sample means ranged from 83.5% capture efficiency in CAN 23 (G 33599) to 91.7% in Santhica 27 (Supplemental Table 3). Six of the 15 individuals had capture efficiency standard errors greater than 1%, Acer II (G 33401), Fasamo (G 33635), Fibridia (G 33623), CAN 23 (G 33599), YunMa (G 33369), and Zaki (G 33214) with YunMa (G 33369) having the largest standard error at 2.8%.

**Figure 4.**
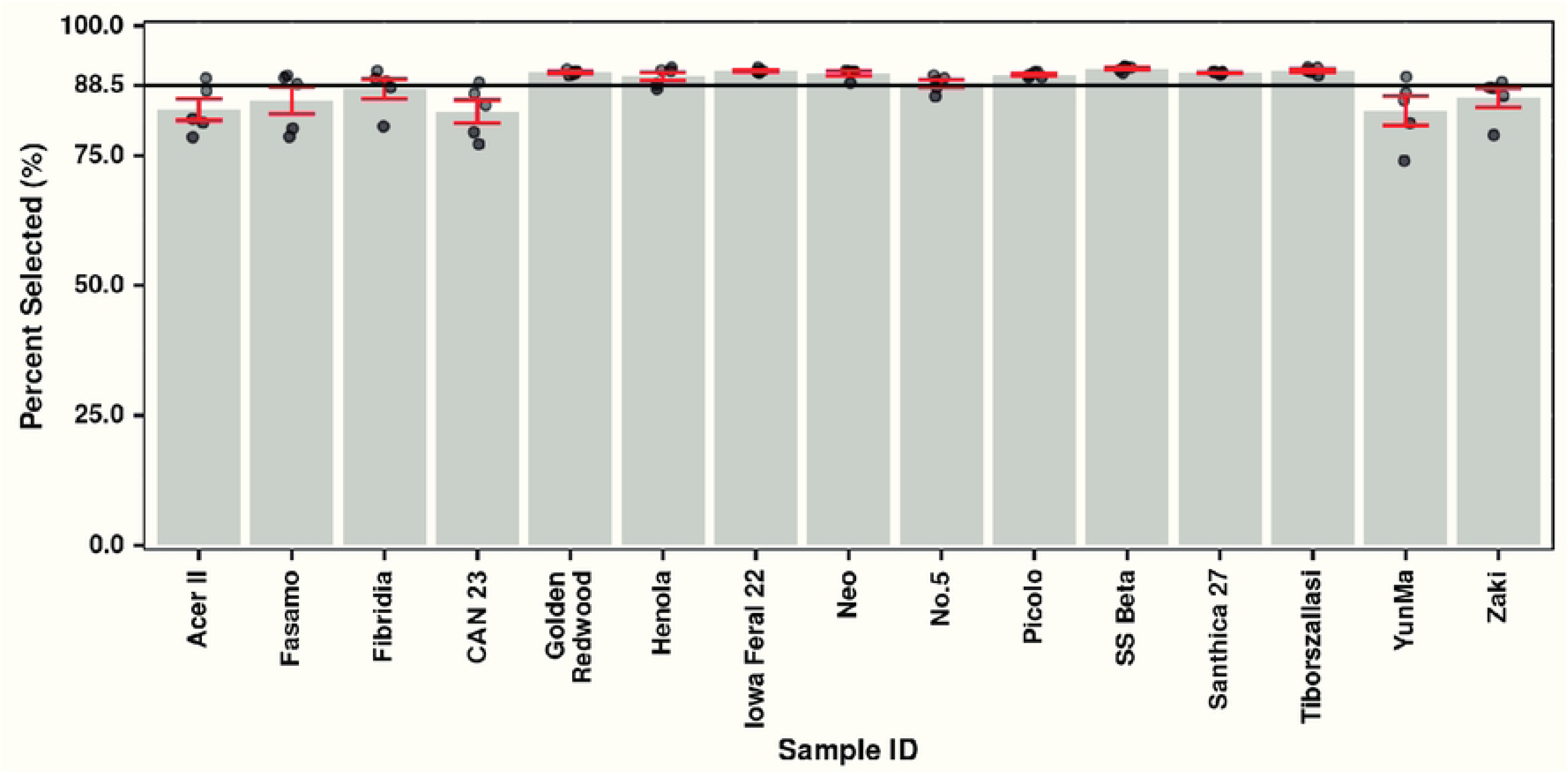
Capture efficiency for 15 replicated individuals genotyped using the CannSelect v1.0 probe set. The solid black line represents the average percent selected across all individuals (88.5%). The gray bars are the mean while the red bars are the standard error of the mean.

### CannSelect v1.0 Probe Alignment Across Diverse Genome Assemblies

The CannSelect v1.0 probe set design was aligned to a diverse set of chromosome-level genome assemblies to show its potential effectiveness in capturing targets across genetic backgrounds. Each assembly had a comparable range of percent total aligned probes, ranging from 94% to 99.8% (Supplemental Table 4). Applying the more stringent specificity filter, the percent of uniquely aligned probes was 94.2% in CBDRx (cs10) and 90.5% in Pink Pepper [18], the current *C. sativa* reference genome (Table 2). Using the diversity groups outlined in Stack et al. [12], the probe set performed best in the high-cannabinoid group (91.1%) while only marginally worse among the lowest group, U.S. Feral (89.3%).

**Table 2:** CannSelect v1.0 Probe Set Alignment and Specificity by Diversity Group.

|  |  | **Aligned (%)** | **Specific (%)** |
| --- | --- | --- | --- |
|  |  | **(Mean ± SE)** | **(Mean ± SE)** |
| **Reference Genomes** | CBDRx (cs10) | 99.8 | 94.2 |
|  | Pink Pepper | 98.5 | 90.5 |
| **Diversity Groups\*** | Asian Hemp (N = 4) | 97.0 ± 0.9 | 89.8 ± 0.9 |
|  | European Hemp (N = 22) | 97.5 ± 0.3 | 90.3 ± 0.3 |
|  | High-Cannabinoid (N = 70) | 98.1 ± 0.1 | 91.1 ± 0.1 |
|  | U.S. Feral (N = 4) | 96.4 ± 1.0 | 89.3 ± 1.0 |

#NCBI RefSeq IDs - CBDRx (cs10): GCF\_900626175.2, Pink Pepper: GCF\_029168945.1;
\*Diversity groups are based on assembly designations used in Stack et al., 2025.

## Discussion

The CannSelect v1.0 probe set is a significant advancement in genotyping technology for *C. sativa* breeding and genomic research. The coverage and probe density achieved by CannSelect v1.0 can provide researchers an effective tool for both trait mapping studies and diversity analysis. Through a GWAS in a previously uncharacterized breeding population, we successfully mapped day-neutral flowering to the *Autoflower1* locus on chromosome 1 [13]. We further demonstrated the capture efficiency of CannSelect v1.0 at close to 90% across diverse hemp germplasm with excellent repeatability. Additionally, *in silico* alignments predict that CannSelect v1.0 will perform well in capturing unique targets across diversity groups in *C. sativa*. The CannSelect probe set offers a path to high-quality sequencing data for *C. sativa* at low cost based on the ability to multiplex reduced representation libraries for sequencing. We expect that the approach used in designing the CannSelect probe set can be effectively followed in the design of similar target-enrichment probe sets for other crop species, accelerating trait discovery and breeding across agricultural systems.

## Materials and Methods

### Probe Design

Target sequences for the CannSelect probe set could be placed into two groups, either marker-based or gene-based. For the marker-based sequences, the targets were built around genome-wide polymorphic sites sourced and filtered from four datasets. Two datasets were downloaded from publicly available repositories, both consisting of whole-genome resequencing data from diversity analyses, Ren et al. [8] and Woods et al. [9]. For the remaining two datasets, the McKay Lab (Colorado State University, CO) provided another set of whole-genome resequencing data of accessions from the Leibniz Institute of Plant Genetics and Crop Plant Research (IPK) in Gatersleben, Germany. Conserved polymorphic SNPs were also extracted from genotyping-by-sequencing (GBS) data from the Cornell hemp breeding program, some of which were described in Carlson et al. as part of a diversity analysis [7].

VCF files were obtained for each dataset, containing roughly 543,000 (Carlson), 1.81 million (McKay IPK), 12.01 million (Ren), and 8.74 million (Woods) markers. All four datasets were aligned to the CBDRx (cs10) [15] genome assembly. Each dataset was filtered independently for 99% call rate, heterozygosity between 5% and 95%, and thinned to a minimum of 140 bp between each marker to avoid overlapping target regions. Filtering was conducted using Tassel 5 [19]. Marker positions from each dataset were then merged and thinned again to prevent overlap. For each marker in the merged dataset, two probes were tiled onto the target polymorphism. This generated two 120 bp probe sequences sharing a 100 bp overlap centered on the marker with 20 bp on either side of the overlap. Probe sequences were extracted from the CBDRx (cs10) genome assembly [15] using the ‘seqinr’ package [20] in R [21].

For the gene-based targets, genes were selected as those involved in abiotic stress tolerance, cannflavin biosynthesis, disease resistance, flowering time, plant size, and trichome density. Special care was taken to include several Y-specific sequences, CBDAS, and other synthases. The initial gene list contained 148 genes. Gene coding sequences were extracted using NCBI [22] from the CBDRx (cs10) genome assembly [15]. As CBDRx lacks a Y chromosome, those sequences were extracted from the JL-Father assembly (NCBI: GCA_013030025.1). Probes designed over coding sequences were tiled similarly to the marker-based probes, however each 120 bp probe shared a 60 bp overlap with its neighbor.

Candidate probe sequences for both marker and gene-based targets were aligned back to the CBDRx (cs10) genome assembly [15] using ‘blastn’ (max_target_seqs 5, evalue 1e-10, perc_identity 30) through the command-line [23]. Due to the file size, probe sequences were queried by chromosome against the full assembly. For CannSelect v0, blast alignments were filtered to those with an alignment length greater than 108 bp (90% of total probe length) and percent identities greater than 90% with no gaps. If there were three or less alignments remaining, the probe was predicted to be specific. After determining that this was inadequate to filter for non-specific probes, we modified the approach for the next version. For CannSelect v1.0, a formula was applied to filter the blast results (score = alignment length - (alignment mismatch + gap size). Probes with three or fewer alignments scoring above 60 were considered specific. Both tiled probes covering a marker-based target must be predicted to be specific for that target to be retained in the design, while 70% of the probes covering a gene-based target needed to be specific for the gene target to be retained.

Following specificity checks, targets were filtered again. This time focusing on genome coverage and to fit under the maximum limit of Agilent’s Tier 1 design. Firstly, marker-based targets in close proximity to gene-based targets (within 250 kb) were removed. Next, marker-based targets were selected to minimize gaps in the design with the intention of thorough coverage of the *C. sativa* genome. This process was completed using custom code written in R [21]. The complete design was then uploaded to Agilent’s online tool SureDesign [16] as a custom SureSelect design. Each probe’s individual count in the design was boosted based on its GC content by SureDesign during the upload process.

### Testing the CannSelect Probe Set

A combined population of 321 segregating genotypes acquired as ‘Carolina Dream’ (N = 274) and ‘BaOx’ (N = 47) were used to test the CannSelect v0 probe set. These seeds were grown in a greenhouse and planted as single plant plots at Cornell AgriTech in Geneva, NY. As a test case for trait mapping, each plant was surveyed for day-neutral flowering in the greenhouse as a binary trait following the methods described in Toth et al. [13]. Leaf samples were collected from each plant and used for library prep, target capture, and sequencing by the Cornell Institute for Biotechnology. Library preparation was completed using the Agilent (Santa Clara, CA) SureSelect XT HS2 DNA Preparation Kit with sequencing performed on an Illumina (San Diego, CA) NextSeq500 (2x150 bp).

Data QC and variant calling were performed on the Cornell BioHPC (https://biohpc.cornell.edu) using a pipeline that closely followed the Genome Analysis Toolkit (GATK) [24] best practices workflow for variant discovery. Raw reads were trimmed using Trimmomatic [25] then fastq files were converted to unmapped bams before alignment to the CBDRx (cs10) [15] genome assembly using BWA-mem [26] under default parameters. At this stage, target-capture data quality and probe specificity were assessed for each sample using ‘CollectHsMetrics’. This output was summarized and visualized using R [21]. Variants were then called for each sample using ‘HaplotypeCaller’, and then jointly across all samples using ‘GenotypeGVCFs’ before hard-filtering based on read depth and quality.

To prepare the data for GWAS, sites were further filtered in R [21] and VCFtools v0.1.16 [27] for biallelic sites that were on-target, with call rates greater than 90%, and minor allele frequencies greater than 0.05. Any missing data from the remaining markers was imputed using LinkImputeR [17]. Principal component and multidimensional scaling analyses were performed on the imputed data using Plink v1.9 [28]. Principal components were saved for use as covariates while population structure was visualized using the MDS components. The day-neutral flowering phenotype was then mapped using a mixed linear model GWAS in Tassel 5 [19] as a binary phenotype with the first five PCs as covariates alongside a kinship matrix to account for population structure. The GWAS results were visualized in R [21].

The CannSelect v1.0 probe set was tested using samples selected to be largely representative of the diversity of the accessions available at the time in the USDA-ARS hemp germplasm repository (Geneva, NY). The samples in this plate included 15 *C. sativa* accessions (Supplemental Table 3), each represented by five random seedling individuals. DNA extraction, hybrid capture, sequencing, and bioinformatics pipeline followed that as described above.

### CannSelect v1.0 Probe Alignment Across Diverse Genome Assemblies

The CannSelect v1.0 probe set was aligned to a diverse set of chromosome-scale genome assemblies using the command-line version of ‘blastn’ (max_target_seqs 5, evalue 1e-10) [23]. In addition to the CBDRx (cs10) [15] and Pink Pepper [18] haploid assemblies, 50 phased diploid genome assemblies from Stack et al. [12] (N = 7), Carey et al. [10] (N = 4), and Lynch et al. [11] (N = 39) were included to capture more genetic diversity. The blast results for each assembly were filtered and checked for specificity using the same method as described above. All filtering and data analysis of blast results were performed in R [21]. Probes were counted as ‘aligned’ if they aligned at least once to the target assembly and ‘specific’ if they had three or less alignments after filtering.

## Acknowledgements

We would like to thank Alex Wares and McKenzie Schessl for excellent technical support and Dr. Qi Sun for bioinformatic guidance. This project was partially funded by Foundation for Food and Agriculture Research (FFAR) grants, NextGen-Hemp-0000000007 with matching funds from Agilent Technologies, and 22-000414, with matching funds from U.S. Sugar. Funding support was also provided by a grant from New York State Department of Agriculture and Markets (CM04068 FR). CAD was supported by an Irish Research Council–Environmental Protection Agency Government of Ireland Postgraduate Scholarship (grant no.: GOIPG/2019/1987).

## Data Availability Statement

Processed genotyping data are available at https://github.com/CornellHemp/CannSelect. The code used in probe design and evaluation are available at https://github.com/wilkersondg/CannSelect, respectively. All other reasonable requests for data should be addressed to the corresponding author.

## Supplemental Information

Supplemental Table 1: CannSelect v1.0 Probe Set Gene-Based Target List.

Supplemental Table 2: CannSelect v0 Design Statistics.

Supplemental Table 3: Capture Efficiency of USDA Accessions.

Supplemental Table 4: Alignment Specificity by Genome Assembly.

